# Wild wood mouse gut microbiota are influenced by captivity and cross-fostering

**DOI:** 10.64898/2026.11.04.764189

**Authors:** Rowan E Bancroft, Sarah E Wolf, Jessica Hall, Amy Sweeny, Amy B. Pedersen

## Abstract

**Background:** Wild and laboratory animals harbour profoundly different gut microbiota, but how microbiomes shift between these states, and the relative roles of captivity, microbial history, and host identity in shaping that shift, remain unclear. Here, we developed a unique wood mouse (*Apodemus sylvaticus*) system comprising free-living mice (Wild:*As*), a wild-derived colony maintained in captivity for over a decade (Wild-like:*As*), and a second colony that we generated from the Wild-like:*As* mice by caesarean rederivation and cross-fostering to laboratory *Mus musculus* (Lab-like:*As*). We compared these three *Apodemus* cohorts with conventional laboratory *Mus musculus* (Lab:*Mm*) to ask how microbial diversity, community composition, and functional potential vary across a spectrum of wild-to-laboratory conditions.

**Results:** We found that alpha diversity varied substantially among cohorts but showed no consistent wild-to-laboratory trend: Shannon diversity was higher in Lab:*Mm* and Lab-like:*As* than in Wild:*As* and Wild-like:*As* mice, whereas phylogenetic diversity was similarly high in both the Wild:*As* and Lab:Mm groups. Community composition more closely reflected cohort histories. Despite over a decade in captivity, Wild-like:*As* mice remained most similar to Wild:*As* conspecifics, whereas Lab-like:*As* mice shifted strongly away from Wild-like:*As* and toward, but remained distinct from, Lab:*Mm* mice. The clearest wild-to-laboratory pattern emerged in microbiome consistency: core ASVs (present in ≥90% of individuals) increased from 9 in Wild:*As* to 46 in Wild-like:*As*, 72 in Lab-like:*As*, and 138 in Lab:*Mm* mice. Predicted functional potential revealed a further distinction between taxonomic and functional patterns: Wild:*As* and Wild-like:*As* microbiomes did not differ significantly, whereas Lab-like:*As* shifted toward the Lab:*Mm* state despite remaining taxonomically distinct.

**Conclusions:** Together, these results show that laboratory-associated microbiomes do not emerge through simple loss of wild microbial diversity, but through broader reorganisation of community composition, consistency, and predicted functional potential reflecting microbial history and host context. The persistent Wild-like and Lab-like microbiome states within *Apodemus* provide a powerful experimental system for future timely work testing how microbiome history influences host physiology, resilience, and responses to environmental challenges.

## Background

Gut microbiota contribute to mammalian physiology and fitness through effects on metabolism, immune function, pathogen resistance, and responses to environmental stress [1–3]. Accordingly, microbiome composition is linked to infection susceptibility, energy balance, and other host phenotypes [4–11]. Microbiome variation can therefore influence experimental outcomes and biological inference. This is particularly relevant for laboratory mice, whose microbial communities differ markedly from those of free-living animals and can be associated with distinct immune and physiological states [12, 13]. These differences raise concerns about how well conventional laboratory mice capture naturally occurring host–microbe interactions.

Laboratory-associated microbiomes often differ from those of free-living animals, and reduced microbial diversity has been reported in laboratory mice and other captive rodents [12, 14, 15]. However, captivity does not produce a uniform response: changes in diet, the environment, social contact, pathogen exclusion, and microbial transmission can reshape gut communities differently across host species and facilities. A meta-analysis across vertebrates found no universal effect of captivity on gut microbial diversity [16], and recent rodent studies reinforce this heterogeneity. Captivity reduced diversity in wild-caught wood mice and Greater white-toothed shrew [17] but altered microbiome composition in pocket mice without detectable changes in alpha diversity [18]. Wild and laboratory mice can also differ in taxonomic composition, predicted functional potential, and temporal dynamics [Hanski et al., 13]. Thus, understanding microbiome change across wild- and laboratory-associated conditions requires moving beyond the expectation of simple diversity loss to ask which dimensions of microbial communities change and whether taxonomic and functional shifts occur in parallel.

Microbiome assembly is also shaped by colonisation history. The order and timing of microbial arrival can generate priority effects, while host genotype and social transmission further constrain which taxa establish and persist [19–22]. These processes are especially relevant to wild to laboratory comparisons because rederivation, cross-fostering, and restricted microbial exposure alter both the microbial pool and routes of acquisition. Wildling mice (laboratory-strain embryos born to wild mice) acquire naturalistic microbiota and immune phenotypes that more closely resemble those of wild mice and humans [23–25], and exposure to wild-derived communities or semi-natural environments can shift conventional laboratory mice toward more wild-associated microbiome and immune states [26–30]. Together, these studies show that microbiome structure reflects not only current housing, but also past microbial exposure and transmission history. Yet these approaches have largely asked what happens when wild-associated microbiota are introduced to laboratory mice; far less is known about how microbiomes of wild-derived hosts are reorganised under laboratory conditions.

The wood mouse (*Apodemus sylvaticus*), a common European rodent with naturally diverse microbial communities [31–33], provides a unique system in which to address this gap. Alongside free-living wood mice, we maintain a wild-derived *A. sylvaticus* colony that has been housed in captivity for more than a decade (Wild:*As* and Wild-like:*As*, respectively), together with a second colony generated from the Wild-like cohort through caesarean rederivation and cross-fostering to laboratory *Mus musculus* (Lab-like:*As*). We compared these three *Apodemus* cohorts with conventional laboratory *Mus musculus* (Lab:*Mm*), creating a series of contrasting microbiome histories spanning wild- to laboratory-associated conditions. We asked how gut microbial diversity, community composition, and predicted functional potential vary across these cohorts, and whether the intermediate *Apodemus* colonies reveal distinct signatures of captivity, microbial history, and host context. This framework allows us to move beyond a simple wild-versus-laboratory comparison and test whether laboratory association is characterised by microbial loss or by broader reorganisation of gut microbial communities.

## Materials and Methods

### Ethics statement

All procedures were conducted under UK Home Office Project Licence 70/8543 in accordance with the Animals (Scientific Procedures) Act 1986 and ScotPIL guidelines.

### Wild to laboratory-reared gradient of cohorts

#### Wild wood mice (Wild:As)

Faecal samples were collected from 27 wild adult wood mice at a Scottish woodland site (Hewan Wood, Polton, Scotland) between July and November, 2020. All samples were collected at first capture prior to any intervention and stored at −80°C within 4–6 h.

#### Wild-like:As wood mouse colony

A captive colony of wild-derived wood mice has been maintained at the University of Edinburgh since 2011. This colony was originally established by J. Clarke in 1995 at the Department of Zoology, University of Oxford, and was maintained with occasional introductions of wild wood mice. The wood mouse colony moved to the University of Liverpool, Faculty of Veterinary Science around 16 years ago [34], and then a subset of the wood mice colony was brought to the University of Edinburgh in 2011. Due to reduced breeding success in this colony over time, in 2016 we brought in a new set of 16 breeding pairs from a long-standing (44 generations) wood mouse colony which was maintained at Fera Science limited. For the past 10 years, we have maintained this new wild-derived wood mouse colony, with between 12-16 outbred male lineages housed under standard specific-pathogen free (SPF) conditions (individually ventilated cages (IVCs), 12 h light cycle, controlled temperature, *ad libitum* food and water, (SDS Rat and Mouse 3). Annual FELASA-based health monitoring is performed (Mähler Convenor et al. 2014). In 2017, PCR screening (IDEXX Bioresearch, Germany) detected *Tritrichomonas muris* and *Spironucleus muris*, and so the colony was housed in a separate facility.

#### Lab-like:As wood mouse colony

To establish a parasite-free colony, Wild-like:*As* embryos were re-derived by caesarean section and cross-fostered to CD1 (*Mus musculus*) dams [Charles River Laboratories;, 35]. Eighteen unrelated Wild-like:*As* breeding pairs were time-matched with mated CD1 females. Caesarean sections were performed at embryonic day 19 under aseptic conditions, and pups were decontaminated in 1% Vikon (Dupont, UK) and retrieved in a sterile B60 Bell Isolator. To reduce maternal rejection, pups were coated with CD1 dam urine prior to fostering. Litters were transferred to CD1 dams (∼4 pups per dam, mixed with CD1 pups) and maintained under SPF conditions in CD1-only housing. At weaning, fostered wood mouse offspring (Lab-like:*As*) were identified by coat colour, and unrelated individuals were paired to establish 6 outbred lineages. The colony was maintained under SPF conditions alongside *Mus musculus* strains in a separate facility from Wild-like:*As* since 2017.

#### CD1 mice (Lab)

Just prior to the onset of sampling, CD1 mice were purchased from Charles River Laboratories (Edinburgh, UK), as the strain was no longer bred in-house.

### Sample collection for gut microbiota characterisation

At the onset of sampling (June - October 2021), subsets of CD1 (n = 36), Lab-like:*As* (n = 36), and Wild-like:*As* (n = 36) mice were transferred into a single shared facility to ensure a common environmental context. All animals were individually identified using ear biopsies and housed in their transport groups in IVCs under identical husbandry conditions, with cohorts maintained on separate racks within the same room.

On the first sampling day, CD1 mice were introduced into the facility first and faecal samples were collected under sterile conditions in a class II laminar flow hood. Only after completion of CD1 sampling were Lab-like:*As* mice introduced, sampled using the same procedure, and assigned their housing, followed sequentially by Wild-like:*As* mice. Faecal sampling was then repeated weekly for 2–3 weeks using an identical protocol. All procedures were performed by the same operator in a fixed order (CD1 → Lab-like:*As* → Wild-like:*As*), and laboratory gloves were changed between handling mouse cohorts to minimise cross-contamination. Cage changes were performed weekly following the same sequence. All animals had *ad libitum* access to autoclaved RM1 chow. Mice were sex-matched where possible and aged 5–13 weeks at first sampling. Faecal samples were stored at −80°C within 4 h of collection.

### DNA extraction and sequencing

DNA was extracted from faecal samples using the QIAamp PowerFecal Pro DNA Kit (Qiagen Ltd, UK) according to manufacturer’s instructions. Extractions were performed in batches of 24 samples, including at least one dH_2_O-only negative control per batch. Briefly, samples were homogenised (1:2 w/v in dH_2_O), and 0.03g aliquots were lysed using PowerBead Pro tubes and lysis buffer. Following lysis, inhibitor removal, binding, and wash steps were performed, and DNA was eluted in 50 µl of elution buffer. DNA yield was assessed using NanoDrop^TM^ spectrophotometry, and absence of amplification was confirmed in negative controls. Extracts were stored at -20°C until further use.

The V4 region of the bacterial 16S rRNA gene was amplified using barcoded primers in a dual-index PCR system [515F/806R; 36]. All PCR set-up was conducted in a UV-sterilised cabinet (SCIE-PLAS Ltd, Cambridge, UK) using UV-treated consumables to minimise contamination. Individual PCR reactions were then set up at a final volume of 50µl using Roche reagents as follows: 37µl Nuclease-free PCR-grade H_2_O, 5µl 10x PCR Buffer, 2µl MgCl_2_ (25 mM), 1µl dNTP mix (10 mM), 0.5µl Taq DNA polymerase (5 U/µl), 1.25µl Forward barcode primer (10µM), 1.25 µl Reverse barcode primer (10 µM), and 1 µl DNA sample. Each sample was assigned a unique primer barcode combination to enable multiplexing.

PCR reactions were performed on 96-well plates containing 91 samples and 5 controls per plate: water-only negative controls, primer-only negative controls, a commercial mock community positive control (ATCC MSA-1003), and two pooled biological positive controls (“Golden Colony” and “Golden Wild”), each generated from pooled DNA of six individual mice. Samples were randomly assigned across plates. The thermocycle consisted of initial denaturation (94°C for 3 minutes), followed by 25 cycles of denaturation (94°C for 45 sec), annealing (50°C for 1 min), and extension (72°C for 1.5 min), with a final extension at 72°C for 10 mins. Amplicon size and lack of contamination were verified using gel electrophoresis.

Amplified DNA was then purified using AMPure XP Beads (Beckman Coulter, UK) and quantified using a Quant-iT PicoGreen dsDNA Assay Kit (Thermo-Fisher, UK). Samples were pooled at equimolar concentrations for sequencing. Negative controls were below detection by PicoGreen but were included in the final pool (2µl of each sample) for downstream sequencing. Sequencing was performed by Edinburgh Genomics, with the addition of custom primers [36] using an Illumina MiSeq v2 platform to generate 250 base pair (bp) paired-end reads and ∼11 million raw reads. Eight sequencing runs were performed, with samples randomly assigned to each run.

### Processing of sequence data

Raw Illumina sequences were processed using the DADA2 pipeline (v1.16) in R [37, 38] with identical parameters across runs. Based on quality profiles, forward and reverse reads were truncated (parameter ‘truncLen’) between 220-240 bp and 170-190 bp, respectively. Reads were filtered using default DADA2 parameters, allowing a maximum of two and five expected errors for forward and reverse reads, respectively, with truncation of reads below a quality score of 2. Amplicon sequence variants (ASVs) were inferred from denoised reads, paired-end reads were merged, and chimeras were removed. Taxonomy was assigned using the DADA2 naive Bayesian classifier against the SILVA database [v138.1; 39]. Mean read retention after processing was ∼80%.

ASVs classified as non-bacterial (Eukaryota, mitochondria, chloroplasts), unassigned taxa, and low-abundance contaminants were removed, resulting in 2,863 unique bacterial ASVs. Potential reagent or laboratory contaminants were identified using a prevalence method that compares ASV occurrence in water-only and water-plus-primer negative controls with biological samples using a classification threshold of 0.5. Seven ASVs were classified as potential contaminants and removed (**Table S1**). Negative controls contained substantially fewer reads than biological samples (median = 138 reads, range = 59–409; **Fig. S1**). Samples with <10,000 reads were excluded (n = 4, mean = 141.5 reads, range 98-222 reads). Mock community and pooled biological (“golden”) controls were included on each sequencing plate to assess sequencing performance and technical reproducibility. The Mock-community profiles (mean = 63,838 reads, range = 33,731 - 71,636 reads) were consistent across plates, as were the golden controls (mean = 67,397 reads, range = 45,063 - 91,124 reads). Thus, a total of 17,550,516 reads (mean = 73,433 reads per sample; range = 16,648 – 737,454 reads), from 239 faecal samples (26 Wild:*As*, 70 Wild-like:*As*, 71 Lab-like:*As* and 72 Lab:*Mm*) were analysed.

### Statistical analysis

#### Alpha Diversity

All statistical analyses were conducted in R version 4.2.1 (R Core Team 2026). Alpha diversity was quantified using Shannon diversity and Faith’s phylogenetic diversity, calculated with the vegan and picante packages, respectively [40, 41]. Shannon diversity incorporates taxonomic richness and evenness [42], whereas Faith’s phylogenetic diversity reflects the total phylogenetic branch length represented within a sample [43]. To standardize sequencing depth, samples were rarefied without replacement to 15,857 reads (the minimum sampling depth) 100 times [44], and the mean diversity value across iterations was used for each sample. Each metric was analysed using a Gaussian GLMM with cohort and sex as fixed effects and mouse ID as a random effect. Overall effects were assessed using Type III Wald χ² tests followed by Tukey-adjusted pairwise comparisons. As a sensitivity analysis, models were repeated using unrarefied diversity estimates. This and analyses of observed richness and inverse Simpson diversity can be found in the Supplemental.

#### Beta Diversity

To evaluate between-sample differences in gut microbial community composition, analyses were first restricted to a single pre-intervention faecal sample per mouse. For lab-reared cohorts, the sample closest to Day 0 was retained when multiple samples were available; one sample per individual was retained for wild mice. ASVs with <5 total reads or occurring in <10% of samples were then removed [45], leaving 595 ASVs for the primary analysis. Read counts were transformed to relative abundances for Bray– Curtis dissimilarity, while Jaccard distance was calculated from ASV presence-absence. Beta diversity was visualised using principal coordinates analysis (PCoA) and assessed using PERMANOVA [adonis2, vegan v2.6-4, 41], with cohort and sex as fixed effects. Pairwise cohort comparisons were subsequently performed with Benjamini-Hichberg correction for multiple testing. Homogeneity of multivariate dispersion was assessed using PERMDISP to determine whether compositional differences were accompanied by differences in within-group heterogeneity. Weighted and unweighted UniFrac were additionally analysed. To assess sensitivity to ASV filtering, Bray-Curtis and Jaccard analyses were repeated across global prevalence thresholds of 0-15%, while retaining the ≥5-read criterion.

#### Differential Abundance

Differential abundance was assessed using MaAsLin3 [46]. As for beta diversity analyses, ASVs with <5 total reads or present in <10% of samples were removed [47]. Counts were converted to total-sum-scaled relative abundances using library sizes calculated before filtering. Complementary genus- and family-level analyses were used to assess broader taxonomic patterns, for which raw ASV counts were aggregated by taxonomic assignment before normalization; only ASVs with informative assignments at the respective rank were included. Genus-assigned ASVs represented 43.6% of ASVs and 72.1% of sequence reads, while family-assigned ASVs represented 79.8% of ASVs and 97.5% of reads. Following filtering, 75 genera and 44 families were retained.

At each taxonomic resolution, cohort, sex, and log10-transformed sequencing depth were included as fixed effects, with individual identity as a random effect. Relative abundance was log-transformed within MaAsLin3, and compositionality was addressed using its median-comparison procedure. All six pairwise comparisons among cohorts were estimated as linear contrasts from the fitted model using maaslin_contrast_test, rather than by fitting separate models for each comparison. Positive coefficients indicate higher abundance in the first-named group. P-values were adjusted using the Benjamini–Hochberg false-discovery rate (FDR), with q < 0.05 considered significant.

#### PICRUSt2 Functional Analysis

Microbial functional potential was inferred using PICRUSt2 [48] and prediction quality was assessed using weighted nearest-sequenced-taxon index (NSTI) scores.

Differences in weighted NSTI among microbiota origins were tested using a linear mixed-effects model with microbiota origin as a fixed effect and mouse identity as a random effect to account for repeated sampling. PICRUsT2-predicted KEGG Ortholog (KO) abundances were aggregated to KEGG pathways using ggpicrust2 [Yang et al., 49]. To reduce sparsity and exclude rarely predicted functions, pathways occurring in <10% of samples were removed, retaining 234 pathways. Robustness to this filtering decision was assessed by repeating the functional beta-diversity analysis across global pathway-prevalence thresholds from 0–20%. Pathway abundances were converted to relative abundances, and Bray-Curtis dissimilarities were used to assess overall functional composition. To avoid pseudoreplication, these analyses used one sample per individual (n = 132), i.e., the sample closest to Day 0 for colony mice and the earliest sample for wild mice. Differences among microbiota origins were visualized using PCoA and tested using PERMANOVA with 9,999 permutations, followed by pairwise PERMANOVAs with Benjamini–Hochberg correction. Homogeneity of multivariate dispersion was assessed using PERMDISP, and the robustness of microbiota-origin effects to sequencing depth was assessed by additionally including log10-transformed read depth. Differential abundance of predicted KEGG pathways was tested using MaAsLin3 with cohort, sex, and log10-transformed sequencing depth as fixed effects and mouse identity as a random effect. All six pairwise contrasts were estimated from the same model, with FDR-adjusted q < 0.05 considered significant. Significant pathways were summarized descriptively by KEGG Level 1 and Level 2 classifications.

## Results

We successfully established and now maintain six genetic lines of Lab-like:As wood mice at the University of Edinburgh, alongside 12 genetic lines of Wild-like:*As*. The Lab-like:*As* colony is housed in the same animal facility as conventional laboratory mice, adjacent to but separate from the Wild-like:*As* colony.

### Overall Composition

Taxonomic composition differed markedly among the four mouse cohorts at both the family and ASV levels (**Fig. 1**). At the family level, cohorts were dominated by members of the Lactobacillaceae, Lachnospiraceae, Muribaculaceae, and Rikenellaceae (**Fig. 1A**). Despite these broad similarities in higher-level taxonomic composition, ASV-level overlap among cohorts was more limited (**Fig. 1B**). The largest proportion of ASVs was unique to the Wild:*As* cohort (972), followed by the Lab:*Mm* cohort (413), whereas comparatively fewer ASVs were unique to the Lab-like:*As* (213) and Wild-like:*As* (147) cohorts. Shared ASVs were most common among the three *Apodemus* cohorts (n = 103). In contrast, the greatest overlap involving Lab:*Mm* occurred with the Lab-like:*As* cohort (192 ASVs), while relatively few ASVs were shared exclusively between Lab:*Mm* and either Wild:*As* (36) or Wild-like:*As* (5). Only 94 ASVs were detected across all four cohorts. These universally shared ASVs were themselves dominated by members of Lachnospiraceae (26 ASVs; 27.7%), Oscillospiraceae (16; 17.0%), and Muribaculaceae (15; 16.0%) families, with smaller contributions from Rikenellaceae (6; 6.4%), Lactobacillaceae (4; 4.3%) and several additional low-abundance families (**Fig. 1B**).

**Figure 1.**
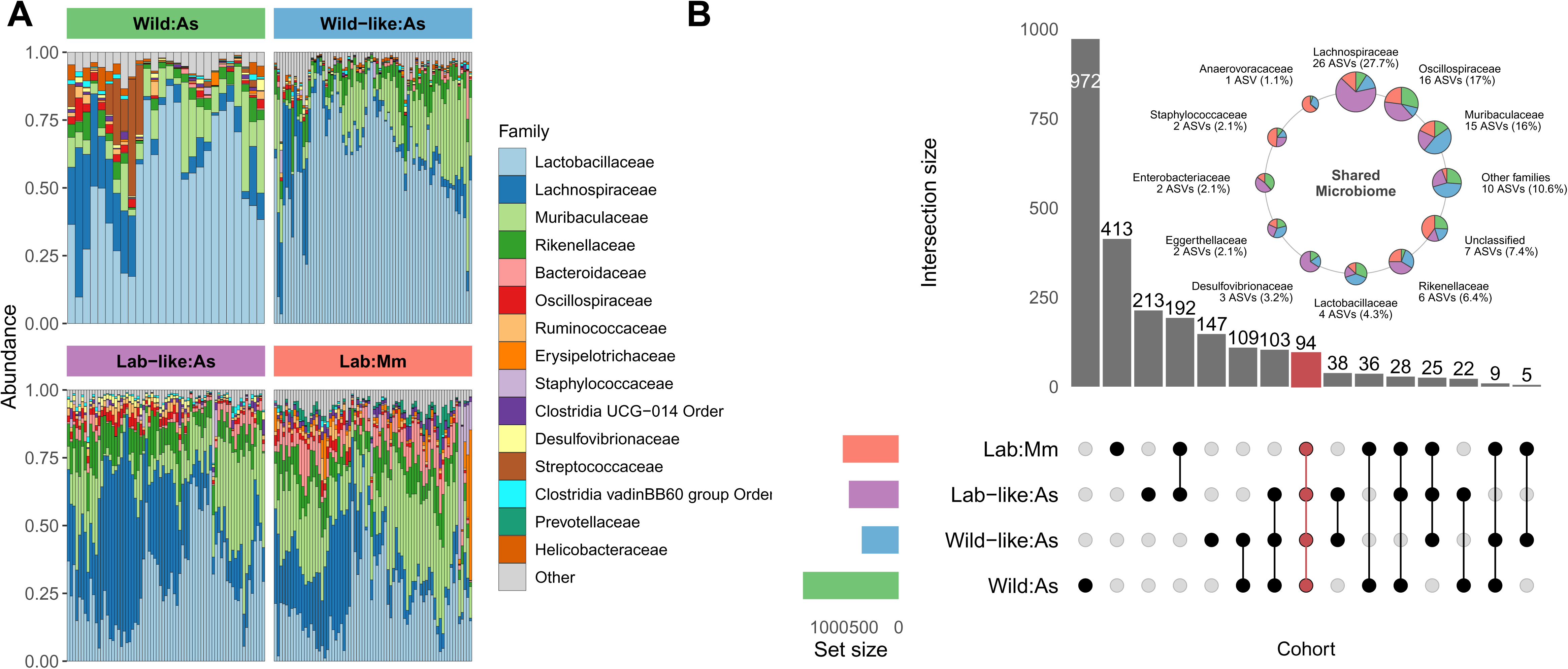
Taxonomic composition and overlap of the gut microbiota across the four mouse cohorts. **(A)** Family-level relative abundance of the gut microbiota in wild-caught wood mice (Wild:*As*, *Apodemus sylvaticus*), wild-derived laboratory-reared wood mice (Wild-like:*As*), caesarean re-derived and laboratory fostered wood mice (Lab-like:*As*), and conventional laboratory mice (Lab:Mm, *Mus musculus*). Each vertical bar represents the microbiota composition of an individual sample. **(B)** UpSet plot showing the overlap of amplicon sequence variants (ASVs) among the four cohorts. Horizontal bars indicate the total number of ASVs detected within each cohort, whereas vertical bars indicate the number of ASVs shared among the cohort combinations indicated by the connected dots below each bar. The red bar and connected red dots highlight the 94 ASVs shared across all four cohorts. The inset summarises this shared microbiome, with each pie chart representing a bacterial family containing one or more of the 94 shared ASVs. Pie size is proportional to the number of shared ASVs assigned to that family, and coloured sectors indicate the mean relative abundance of that family within each cohort.

### Core Microbiome

The size of the core microbiome differed markedly among the four cohorts. Using a core definition of ASVs present in ≥90% of individuals within a cohort, Wild:*As* had only 9 core ASVs, compared with 46 in Wild-like:*As*, 72 in Lab-like:*As*, and 138 in Lab:*Mm*. Core membership was also largely cohort-specific: of the 214 ASVs identified as core in at least one cohort, only three were core across all four cohorts, whereas 170 were core in only a single cohort. Lab:*Mm* contained the greatest number of cohort-specific core ASVs (104), followed by Lab-like:*As* (37), Wild-like:*As* (27), and Wild:*As* (2). This pattern was robust to the prevalence threshold used to define the core microbiome: from 50% to 100% prevalence, Lab:*Mm* consistently had the largest core and Wild:*As* the smallest, with the two wild-derived colonies at intermediate levels.

### Alpha Diversity

Gut microbial alpha diversity differed strongly among cohorts for both Shannon diversity (Wald χ²□= 167.24, *P* < 0.001) and Faith’s phylogenetic diversity (χ²□= 161.92, *P* < 0.001), whereas sex was not associated with either metric (Shannon: *P* = 0.727; Faith’s PD: *P* = 0.597). Shannon diversity was higher in Lab:*Mm* and Lab-like:*As* than in Wild-like:*As* (both *P* < 0.001) and Wild:*As* (both *P* < 0.001), while Lab:*Mm* and Lab-like:*As* did not differ (*P* = 0.157) and Wild-like:*As* and Wild:*As* mice were not significantly different (*P* = 0.061; **Fig. 2A**). In contrast, Faith’s PD was similar between Lab:*Mm* and Wild:*As* (*P* = 0.857) and between Lab-like:*As* and Wild-like:*As* (*P* = 0.893), with both Lab-like:*As* and Wild-like:*As* mice exhibiting lower phylogenetic diversity than the Lab:*As* and Wild:*As* cohorts (all *P* < 0.001; **Fig. 2B**). Results were qualitatively consistent for observed richness and inverse Simpson diversity, and conclusions were robust to analyses of unrarefied data (**Fig. S2-S3; Tables S2-S3**).

**Figure 2.**
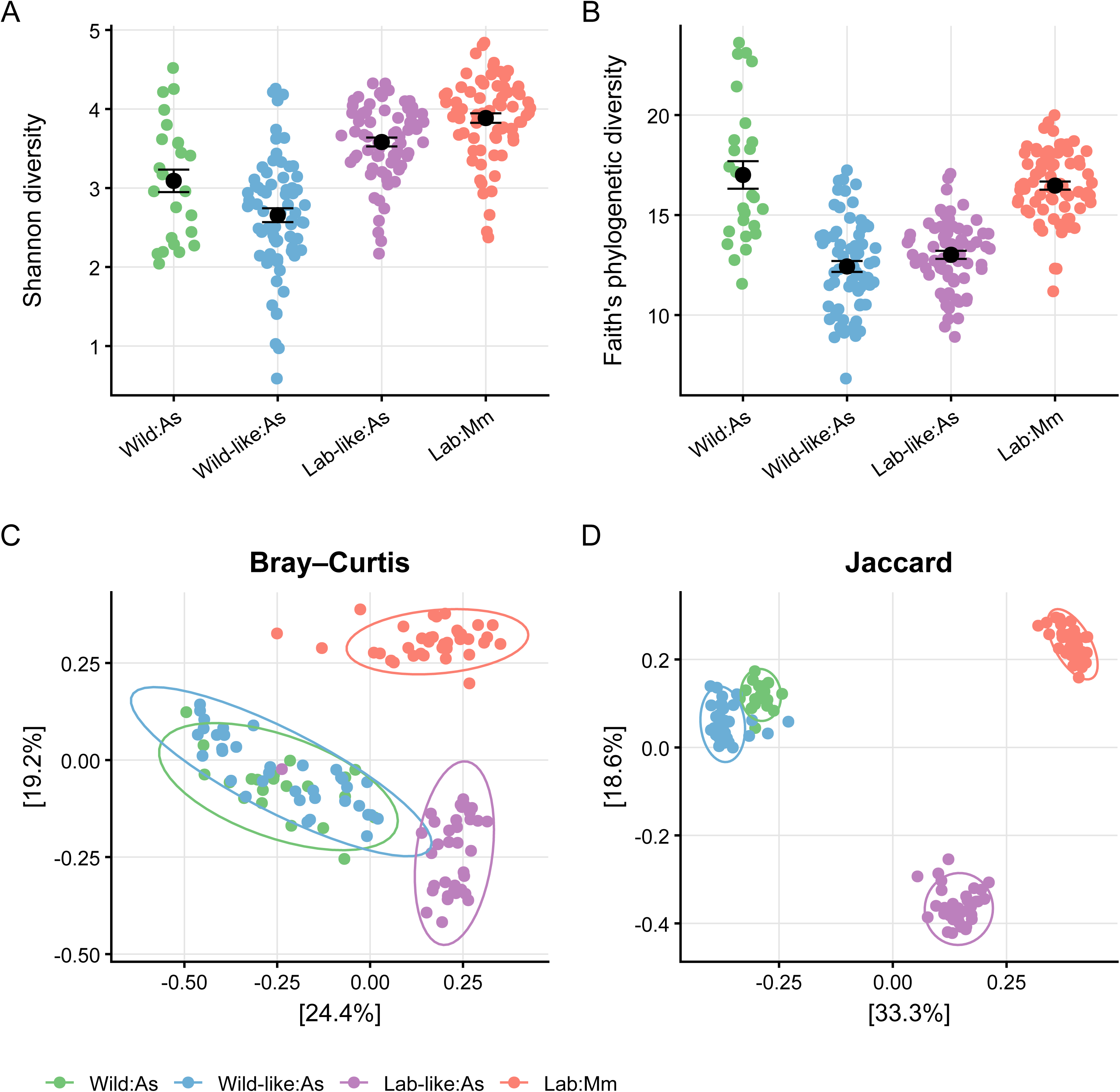
Gut microbiota alpha and beta diversity differ among mouse cohorts. Alpha and beta diversity of 16S rRNA gene sequences from faecal samples collected from four mouse cohorts: wild *Apodemus sylvaticus* (Wild:*As*), wild-derived, laboratory-reared *A. sylvaticus* (Wild-like:*As*), caesarean re-derived and laboratory mouse-fostered *A. sylvaticus* (Lab-like:*As*), and conventional laboratory *Mus musculus* (Lab:Mm). **(A)** Shannon diversity and **(B)** Faith’s phylogenetic diversity. Points represent individual samples, and error bars indicate ± SE of the cohort mean. Higher values indicate greater within-sample microbial diversity. **(C)** Bray–Curtis and **(D)** Jaccard principal coordinates analyses (PCoA) of gut microbial community composition. Each point is an individual mouse. Ellipses indicate the 95% confidence interval per group.

### Beta Diversity

Gut microbial community composition differed markedly among the four mouse cohorts (**Fig. 2C-D**). Cohort explained 41.5% of variation in Bray-Curtis dissimilarity (pseudo- *F□*,□□□= 29.54, *R*² = 0.415, *P* < 0.001) and 56.6% of variation in Jaccard distance (pseudo-*F□*,□□□= 54.55, *R*² = 0.566, *P* < 0.001, **Table S4, Fig. S4**). Pairwise comparisons were significant between every cohort pair for both distance measures (all BH-adjusted *P* < 0.001). For Bray–Curtis dissimilarity, Wild:*As* and Wild-like:*As* were the most similar cohorts (*R*² = 0.141), whereas contrasts among the remaining cohorts showed substantially greater separation (*R*² = 0.315–0.366), with the largest difference between Wild:*As* and Lab:*Mm* (*R*² = 0.366). Presence-absence composition showed a similar pattern: Wild:*As* and Wild-like:*As* again exhibited the smallest difference (*R*² = 0.198), whereas the greatest separation occurred between Wild-like:*As* and Lab:*Mm* (*R*² = 0.564, **Fig. 2D; Table S5**). Multivariate dispersion did not differ among cohorts for Bray-Curtis (*P* = 0.261), whereas dispersion differed significantly for Jaccard (*P* < 0.001), indicating greater heterogeneity in presence–absence composition (**Table S6**). Results were robust to ASV-filtering stringency: cohorts effects remained significant across global prevalence thresholds from 0-15% (**Fig. S5**).

### Differential Abundance

Microbial abundance differed substantially among sample origins. At the ASV level, 61 ASVs differed in at least one pairwise comparison (FDR q < 0.05). Wild-like:*As* and Wild:*As* were the least differentiated cohorts (16 ASVs), whereas 22–32 ASVs differed in all other pairwise comparisons (**Fig. 3**). Cohort differences persisted at broader taxonomic levels. Thirty-eight of 75 genera differed among cohorts. Wild-like:*As* and Wild:*As* appeared the least different (12 genera) compared with 17–22 genera for other comparisons. Several genera consistently differed by cohort, with Ligilactobacillus, Rikenella, Rikenellaceae RC9 gut group, Bacteroides, and Parasutterella differing in five of six comparisons. The direction of these differences varied among cohort contrasts, indicating that laboratory association did not produce a uniform shift in taxon abundance. At family level, 19 families differed among cohorts. Only four distinguished Wild:*As* from Wild-like:*As*, whereas 8–12 differed in the remaining comparisons.

**Figure 3.**
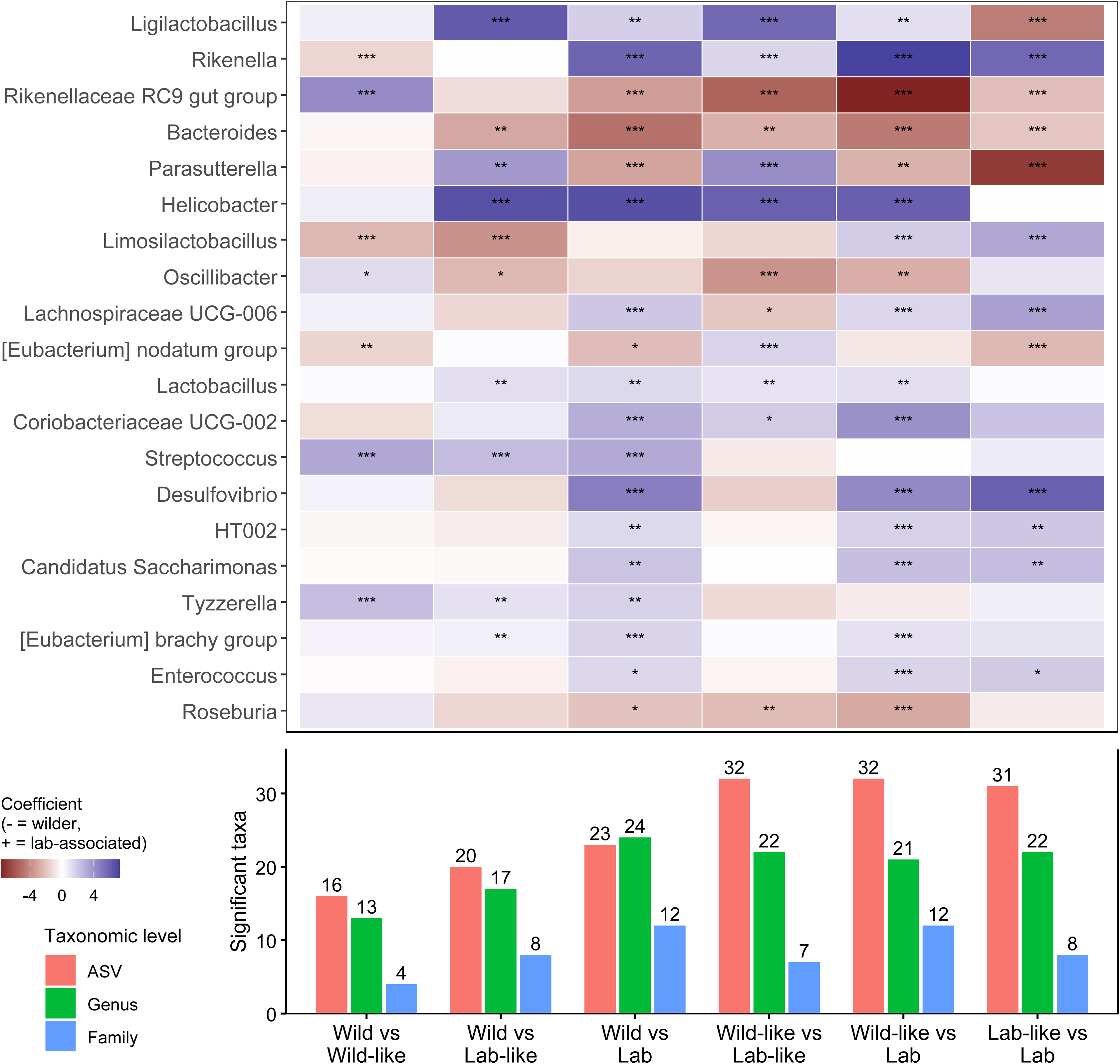
Differential abundance of gut microbial taxa among cohorts. The heatmap shows MaAsLin3 abundance coefficients for the 20 genera showing the strongest and most recurrent differences across pairwise comparisons. Positive coefficients indicate higher abundance in the first-listed cohort and negative coefficients higher abundance in the second; asterisks indicate significant associations (FDR *q* < 0.05). The bar plot below shows the total number of significantly different taxa for each comparison at ASV, genus, and family levels.

### Functional Analysis

PICRUSt2 prediction quality was comparable among cohorts (mean weighted NSTI = 0.045–0.048; LMM: *F□*,□□□= 1.84, *P* = 0.144, **Table S7**). Predicted KEGG composition differed strongly among cohorts (PERMANOVA: *F□*,□□□= 16.12, *R*² = 0.274, *P* < 0.001; **Fig. 4A**), with PCoA1 and PCoA2 explaining 58.5% and 21.2% of variation, respectively. This effect remained strong after accounting for sequencing depth (marginal PERMANOVA: SampleOrigin, *F□*,□□□= 16.09, R² = 0.269, P < 0.001), although sequencing depth explained a small additional proportion of variation (*F□*,□□□= 3.27, R² = 0.018, P = 0.034). The result was also insensitive to pathway-prevalence filtering: although increasing the prevalence threshold from 0% to 20% progressively reduced the number of retained pathways, the PERMANOVA effect size remained effectively unchanged (**Fig. S6**). Multivariate dispersion did not differ among cohorts (*F□*,□□□= 1.84, *P* = 0.141). Pairwise differences were weakest between Wild:*As* and Wild-like:*As* cohorts (*R*² = 0.023, *P* = 0.229) and strongest between Wild-like:*As* and Lab-like:*As* cohorts (*R*² = 0.288, *P* < 0.001; **Fig. 4B**). Lab-like:*As* and Lab:*Mm* composition also remained significantly differentiated (*R*² = 0.059, *P* = 0.010).

**Figure 4.**
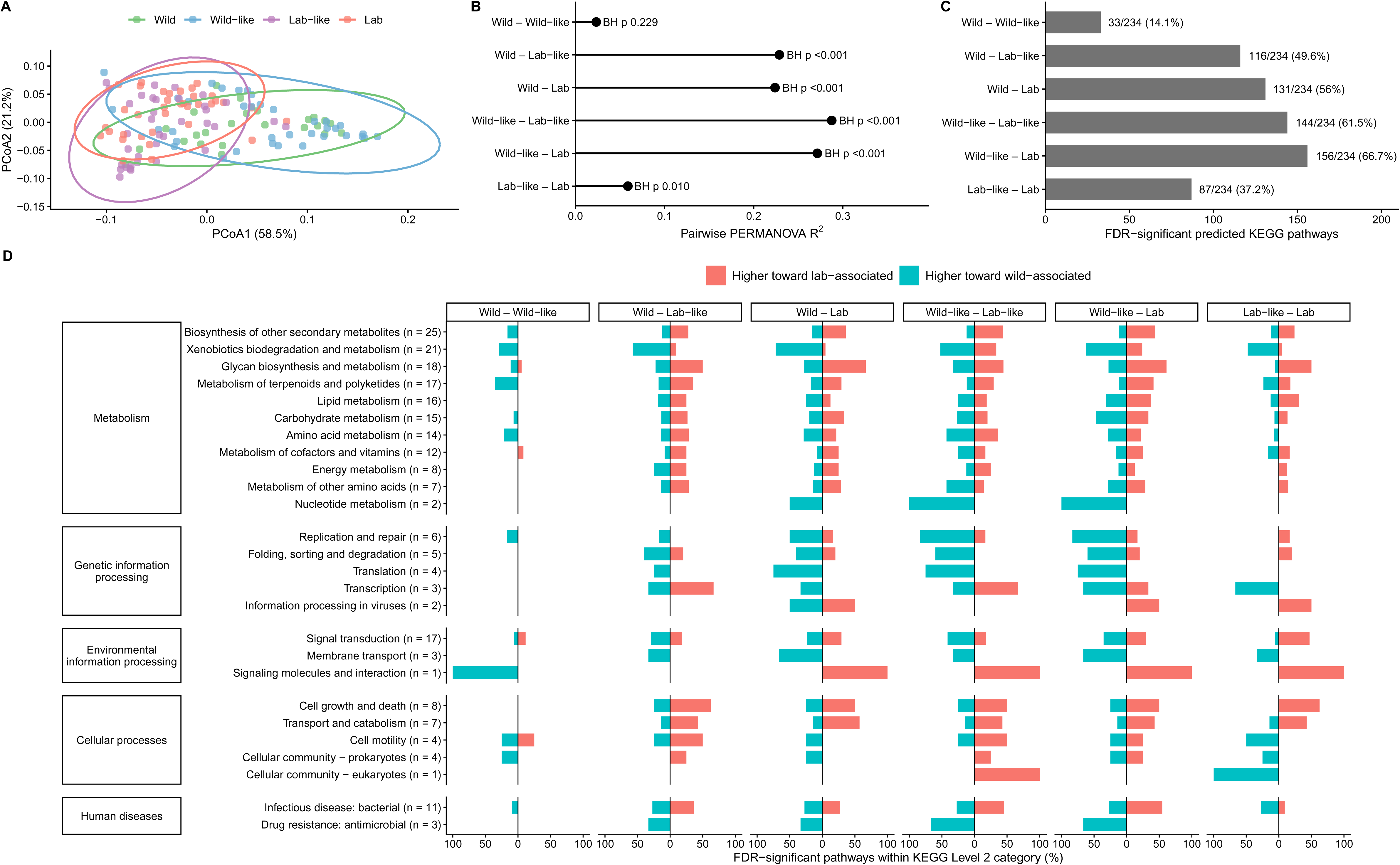
Predicted microbial functional potential across microbiota origins. **(A)** Principal coordinates analysis (PCoA) of Bray–Curtis dissimilarities in PICRUSt2-inferred KEGG pathway profiles; ellipses indicate 95% confidence regions for each microbiota origin. **(B)** Pairwise PERMANOVA effect sizes (R^2^) for differences in predicted functional composition. **(C)** Number of predicted KEGG pathways differing significantly between each pair of microbiota origins (MaAsLin3, FDR q<0.05q<0.05); labels show the number significant out of 234 pathways tested and the corresponding percentage. **(D)** Percentage of pathways within each KEGG Level 2 category that differed significantly between cohorts, with direction indicating higher predicted abundance toward the more wild-associated (teal) or laboratory-associated (coral) microbiota. Numbers in parentheses indicate the total number of pathways tested within each Level 2 category, with categories grouped by KEGG Level 1 classification.

Pathway-level differences showed the same pattern (**Fig. 4C**): only 32/234 pathways (13.7%) differed between Wild:*As* and Wild-like:*As* cohorts, compared with 144/234 (61.5%) between Wild-like:*As* and Lab-like:*As* and 153/234 (65.4%) between Wild-like:*As* and Lab:*Mm*; 89/234 (38.0%) differed between Lab-like:*As* and Lab:*Mm*. Differential pathways were distributed across multiple KEGG functional categories and frequently differed in both directions within the same category (**Fig. 4D**), indicating broad functional reorganization rather than uniform shifts in specific functional classes.

## Discussion

Gut microbiome structure across wild- and laboratory-associated mice reflected both host species (*Apodemus sylvaticus -* As or *Mus musculus -* Mm) and microbial exposure. Despite more than a decade in captivity, Wild-like:*As* remained most similar to free-living Wild:*As* conspecifics. By contrast, caesarean rederivation and cross-fostering shifted Lab-like:*As* microbiomes strongly away from the Wild-like:*As* state and toward Lab:*Mm*, although they remained taxonomically distinct from Lab:*Mm* mice. Predicted functional profiles converged more strongly than taxonomy: Wild:*As* and Wild-like:*As* remained similar, whereas Lab-like:*As* shifted toward the Lab:*Mm* functional state despite retaining a distinct taxonomic composition. Alpha diversity did not decline consistently from wild to laboratory conditions; instead, the clearest directional pattern was increasing microbiome consistency among individuals, with progressively larger core communities from Wild:*As* to Lab:*Mm*. Together, these results show that laboratory-associated microbiomes are not simply depleted versions of wild communities; they are alternative community states shaped by both past microbial exposure and the biological constraints of the host.

The persistence of a recognisably wild-associated microbiome in the Wild-like:*As* colony is particularly striking. Wild-derived microbiota can persist for years under laboratory conditions [21, 50, 51], and we extend this pattern well beyond a decade: Wild-like:*As* remained most similar to wild conspecifics despite prolonged laboratory exposure and repeated opportunities for microbial replacement. One explanation is continued maternal transmission, repeatedly seeding offspring with a similar wild-associated microbial pool. In wild wood mice, maternal transmission dominates early in life before social and environmental transmission becomes increasingly important [22, 52]. By contrast, Lab-like:*As* founders were delivered by caesarean section and fostered by laboratory *M. musculus* dams, replacing this early source with a laboratory-associated microbial pool. Because early colonisation can generate persistent priority effects [20, 53, 54], rederivation and cross-fostering likely helped establish a distinct Lab-like:*As* microbiome within the same wild-derived species.

Lab-like:*As* mice nevertheless remained taxonomically distinct from Lab:*Mm*, indicating that altered early microbial exposure did not override host filtering. Host physiology, immunity, and genetics can favour particular microbial lineages [55–57]. Transplantation experiments further show that introduced microbiota are restructured toward the recipient host, while native bacterial lineages can outcompete non-native ones [58, Sprockett et al., 59]. Host genetic effects also persist within species, including in humans and wild-derived mice [21, 60–62]. Consistent with this, rederivation and cross-fostering shifted Lab-like:*As* strongly toward the laboratory state without erasing a distinct *Apodemus* taxonomic signature. Strikingly, this taxonomic separation was much greater than the predicted functional difference between Lab-like:*As* and Lab:*Mm*. Functional redundancy provides a plausible explanation: different taxa can encode and express overlapping functions [63, 64]. Thus, the *Apodemus* host may preserve a distinct taxonomic community even as laboratory-associated functional potential converges.

Wild mice experience continually changing microbial inputs through diet, season, social contact, and the environment, repeatedly restructuring gut communities [65–67]. Greater exposure in the wild may increase the pool of taxa encountered without uniformly increasing alpha diversity: transient dominance can lower Shannon diversity, while diverse microbial inputs may maintain broad phylogenetic breadth. Consistent with this, Wild:*As* mice had lower Shannon diversity than Lab:*Mm* but similarly high phylogenetic diversity, suggesting less even communities spanning a broad range of bacterial lineages. Both captive *Apodemus* colonies, however, showed lower phylogenetic diversity, consistent with filtering or loss of evolutionary lineages under captivity. The clearest wild-to-laboratory pattern, however, was in between-host consistency: core ASVs (present in ≥90% of individuals) increased from 9 in Wild:*As* to 46 in Wild-like:*As*, 72 in Lab-like:*As*, and 138 in Lab:*Mm*. This mirrors evidence that wild mouse microbiomes are more dynamic and individualised than laboratory communities [Hanski et al., 13, 66, 68]. Standardised diets and restricted microbial sources may instead favour more repeatable communities in captivity. Laboratory association therefore may reorganise both within-host diversity and among-host consistency.

Taxonomic composition showed an even clearer signature of cohort history than alpha diversity. Wild:*As* and Wild-like:*As* were the most similar cohorts in beta diversity and were also the least differentiated at ASV, genus, and family levels. By contrast, comparisons involving laboratory-associated cohorts showed substantially broader taxonomic restructuring. Several recurrent genera reflected this shift: *Ligilactobacillus*, *Rikenella*, *Helicobacter*, and *Streptococcus* tended to be associated with the Wild/Wild-like state, whereas *Bacteroides* and Rikenellaceae RC9 gut group were more prominent toward laboratory-associated states. However, individual taxa did not shift uniformly across cohorts, suggesting that laboratory association reorganises particular bacterial lineages rather than producing a single directional replacement of a “wild” microbiome. The repeated similarity of Wild:*As* and Wild-like:*As* across scales provides especially strong evidence that a recognisably wild-associated community structure persisted after long-term captivity.

Predicted functional profiles also differed across cohorts: Wild:*As* and Wild-like:*As* were similar, whereas Lab-like:*As* shifted toward Lab:*Mm* despite remaining taxonomically distinct, consistent with functional redundancy. KEGG Level 2 differences spanned many categories, indicating broad reorganisation of metabolic potential. Wild-associated profiles showed greater predicted xenobiotic biodegradation and metabolism, consistent with seasonally variable diets and chemically diverse substrates in free-living rodents [65, 66, 69]. Replication and repair, protein folding and degradation, and translation also tended to be higher in wild-associated profiles, potentially reflecting greater microbial turnover or responses to environmental heterogeneity. Laboratory-associated profiles instead shifted across glycan, carbohydrate, lipid, and other metabolic pathways, consistent with a different metabolic organisation under stable conditions. At finer resolution, pathways within the same KEGG category often shifted in opposite directions, indicating reorganisation of specific biochemical routes rather than whole functional systems.

## Conclusion

Rather than forming a simple continuum from a diverse wild microbiome to an impoverished laboratory state, our cohorts represent distinct community states shaped by microbial history, host filtering, and environment. The persistence of contrasting Wild-like and Lab-like microbiomes within *A. sylvaticus* makes this a particularly tractable system for comparing alternative taxonomic and functional states within the same wild-derived host species. Future perturbation experiments can therefore move beyond describing wild-laboratory differences to test whether microbiome history shapes host physiology, resilience, and ultimately fitness.

## Declarations

### Consent for publication

Not applicable.

### Availability of data and materials

The data and code used to reproduce the analyses and figures in this study are available on GitHub at https://github.com/arsweeny/LabToWildMicrobiome. Raw sequencing data is available through the European Nucleotide Archive (Accession PRJEB129335).

### Author Contributions

A.B.P., A.S., and R.E.B. conceptualized and designed the study. R.E.B. and J.H. performed the experiment and collected data. S.E.W., A.S., and R.E.B. analyzed the dataset and prepared all figures. S.E.W. and R.E.B. wrote the main manuscript. All authors reviewed the manuscript.

## Acknowledgements

We thank the Biological and Veterinary Sciences (BVS) team at the University of Edinburgh, especially Gidona Goodman, Craig Watt, Laraine Wells, Richard Trojca, Jenni Russell, David McIver, and David MacDonald, for their invaluable support with maintenance of the wood mouse colonies. We would like to thank Hannah Lemon for assistance with the DNA extractions and sequencing preparation. We also thank Agata Delnicka, Sam Hillman, Julia MacIver and Madeline Hodgeman for their assistance with the collection of wild wood mouse samples in the field.

