## Supplemental for "Wild wood mouse gut microbiota are influenced by captivity and cross-fostering"

**Supplemental Material for “Wild wood mouse gut microbiota are influenced by captivity and cross-fostering”**

| **Metric** | **Title** | **Page** |
| --- | --- | --- |
| QC | Figure S1. Sequencing depth and controls. | 2 |
| QC | Table S1. Contaminant sequence information | 3 |
| Alpha | Figure S2. Other gut microbial alpha diversity metrics among cohorts. | 5 |
| Alpha | Figure S3. Sensitivity of alpha-diversity results to rarefaction. | 6 |
| Alpha | Table S2. Model results for four alpha diversity metrics | 6 |
| Alpha | Table S3. Pairwise comparisons of alpha diversity among cohorts. | 7 |
| Beta | Figure S4. Beta-diversity among cohorts across four distance metrics. | 8 |
| Beta | Figure S5. Sensitivity of beta-diversity results to prevalence filtering. | 9 |
| Beta | Table S4. Overall PERMANOVA results for beta diversity. | 10 |
| Beta | Table S5. Pairwise PERMANOVA results. | 11 |
| Beta | Table S6. Homogeneity of multivariate dispersion among cohorts. | 12 |
| KEGG | Figure S6. Sensitivity of KEGG to pathway-prevalence filtering. | 12 |
| KEGG | Table S7. Weighted NSTI values for PICRUSt2 predictions. | 13 |

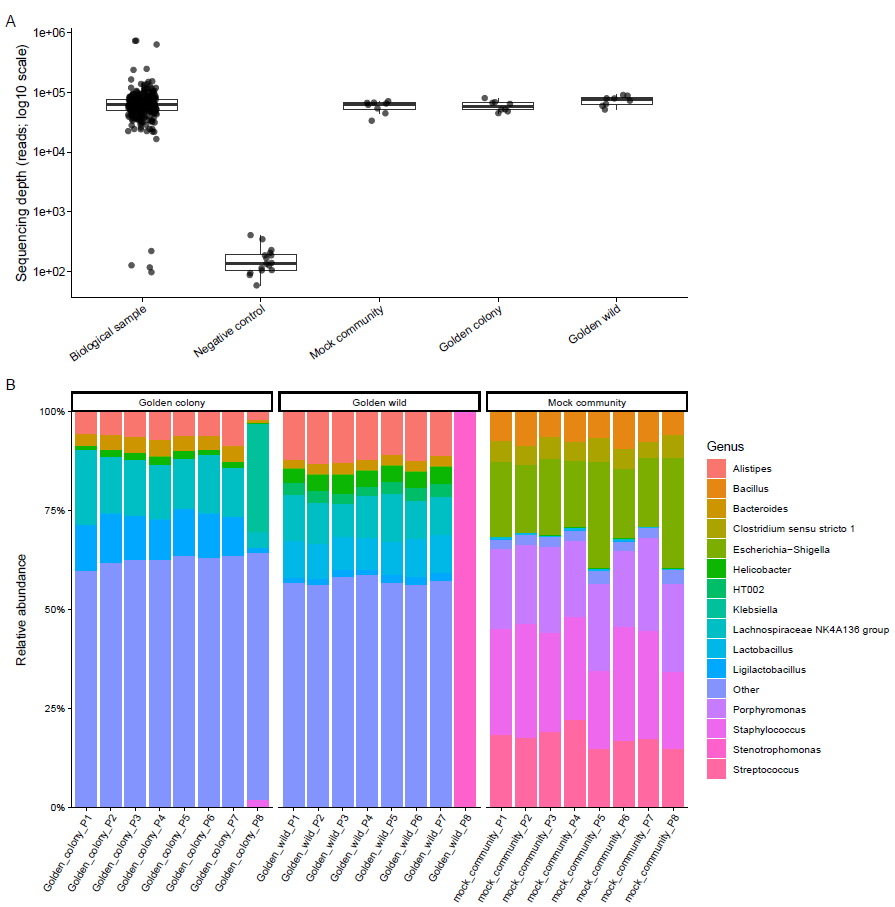

**Figure S1. Sequencing depth and controls. (A)** Sequencing depth for biological faecal samples, negative controls (water-only and water-plus-primer controls), mock-community controls, and pooled biological (“golden”) colony and wild reference samples. Points represent individual samples and boxplots show the median and interquartile range; read depth is displayed on a log10 scale. **(B)** Genus-level relative abundance profiles for repeated golden colony, golden wild, and mock-community controls included across eight sequencing plates. The 15 most abundant genera are shown individually, with remaining genera grouped as “Other”; each stacked bar represents one control sample.

| **Table S1. Putative contaminant amplicon sequence variants (ASVs) detected in negative controls.** ASV sequences, read counts in negative controls and biological samples, and corresponding taxonomic classifications are presented for ASVs identified as potential contaminants. Taxonomic assignments are reported from kingdom to genus where available. | | | |
| --- | --- | --- | --- |
| **ASV Sequence** | **Reads in Negative Controls** | **Reads in Biological Samples** | **Kingdom / Phylum / Class / Order / Family / Genus** |
| TACGTAGGTGGCAAGCGTTGTCCGGATTTACTGGGCGTAAAGGGAGCGTAGGTGGATATTTAAGTGGGATGTGAAATACTCGGGCTTAACCTGGGTGCTGCATTCCAAACTGGATATCTAGAGTGCAGGAGAGGAAAGTAGAATTCCTAGTGTAGCGGTGAAATGCGTAGAGATTAGGAAGAATACCAGTGGCGAAGGCGACTTTCTGGACTGTAACTGACACTGAGGCTCGAAAGCGTGGGGAGCAAACAGG | 5 | 6 | Firmicutes, Clostridia, Clostridiales, Clostridiaceae, Clostridium sensu stricto 1 |
| TACGAAGGGTGCAAGCGTTACTCGGAATTACTGGGCGTAAAGCGTGCGTAGGTGGTCGTTTAAGTCCGTTGTGAAAGCCCTGGGCTCAACCTGGGAACTGCAGTGGATACTGGGCGACTAGAGTGTGGTAGAGGGTAGCGGAATTCCTGGTGTAGCAGTGAAATGCGTAGAGATCAGGAGGAACATCCATGGCGAAGGCAGCTACCTGGACCAACACTGACACTGAGGCACGAAAGCGTGGGGAGCAAACAGG | 47 | 973 | Proteobacteria, Gammaproteobacteria, Xanthomonadales, Xanthomonadaceae, Stenotrophomonas |
| TACGTAGGTGGCAAGCGTTATCCGGAATTATTGGGCGTAAAGCGCGCGTAGGCGGTTTTTTAAGTCTGATGTGAAAGCCCACGGCTCAACCGTGGAGGGTCATTGGAAACTGGAAAACTTGAGTGCAGAAGAGGAAAGTGGAATTCCATGTGTAGCG  GTGAAATGCGCAGAGATATGGAGGAACACCAGTGGCGAAGGCGACTTTCTGGTCTGTAACTGACGCTGATGTGCGAAAGCGTGGGGATCAAACAGG | 16 | 99 | Firmicutes, Bacilli, Staphylococcales, Staphylococcaceae, Staphylococcus |
| TACGGAGGGCGCAAGCGTTAATCGGAATCACTGGGCGTAAAGCGCGCGTAGGCTGTTGTGTAAGTCAGGGGTGAAAGCCCGCGGCTCAACCGCGGAATTGCCTTTGAAACTGCGCAACTTGAATCCAGGAGAGGGCGGCGGAATTCCAGG  TGTAGGAGTGAAATCCGTAGATATCTGGAGGAACACCAGCGGCGAAGGCGGCCGCCTGGCCTGGCATTGACGCTGAGGCGCGAAAGCGTGGGGAGCAAACAGG | 3 | 845 | Desulfobacterota, Desulfovibrionia, Desulfovibrionales, Desulfovibrionaceae, Desulfovibrio |
| TACGTAGGGGGCAAGCGTTATCCGGATTTACTGGGTGTAAAGGGAGCGTAGACGGCAGCGCAAGTCTGAAGTGAAATGCCAGGGCTTAACCCTGGAACTGCTTTGGAAACTGTGCAGCTAGAGTGCAGGAGAGGTAAGTGGAATTTCTAGTGTAGCGGTGAAATGCGTAGATATTAGGAGGAACACCAGTGGCGAAGGCGGCTTACTGGACTGTAACTGACGTTGAGGCTCGAAAGCGTGGGGAGCAAACAGG | 51 | 22585 | Firmicutes, Clostridia, Lachnospirales, Lachnospiraceae, Lachnospiraceae NK4A136 group |
| TACGTAGGTGGCAAGCGTTGTCCGGATTTATTGGGCGTAAAGGGAGCGCAGGTGGTTTCTTAAGTCTGATGTGAAAGCCCACGGCTTAACCGTGGAGGGTCATTGGAAACTGGGAAACTTGAGTACAGAAGAGGAATGTGGAACTCCATGTGTAG  CGGTGGAATGCGTAGATATATGGAAGAACACCAGTGGCGAAGGCGACATTCTGGTCTGTTACTGACACTGAGGCTCGAAAGCGTGGGGAGCAAACAGG | 6 | 2730 | Firmicutes, Bacilli, Lactobacillales, Aerococcaceae, Aerococcus |
| TACGTAGGTGGCAAGCGTTGTCCGGATTTATTGGGCGTAAAGCGAGCGCAGGCGGAAGAATAAGTCTGATGTGAAAGCCCTCGGCTTAACCGAGGAACTGCATCGGAAACTGTTTTTCTTGAGTGCAGAAGAGGAGAGTGGAACTCCATGTGTAGCGGTGGAATGCGTAGATATATGGAAGAACACCAGTGGCGAAGGCGGCTCTCTGGTCTGCAACTGACGCTGAGGCTCGAAAGCATGGGTAGCGAACAGG | 8 | 11572 | Firmicutes, Bacilli, Lactobacillales, Lactobacillaceae, Lactobacillus |

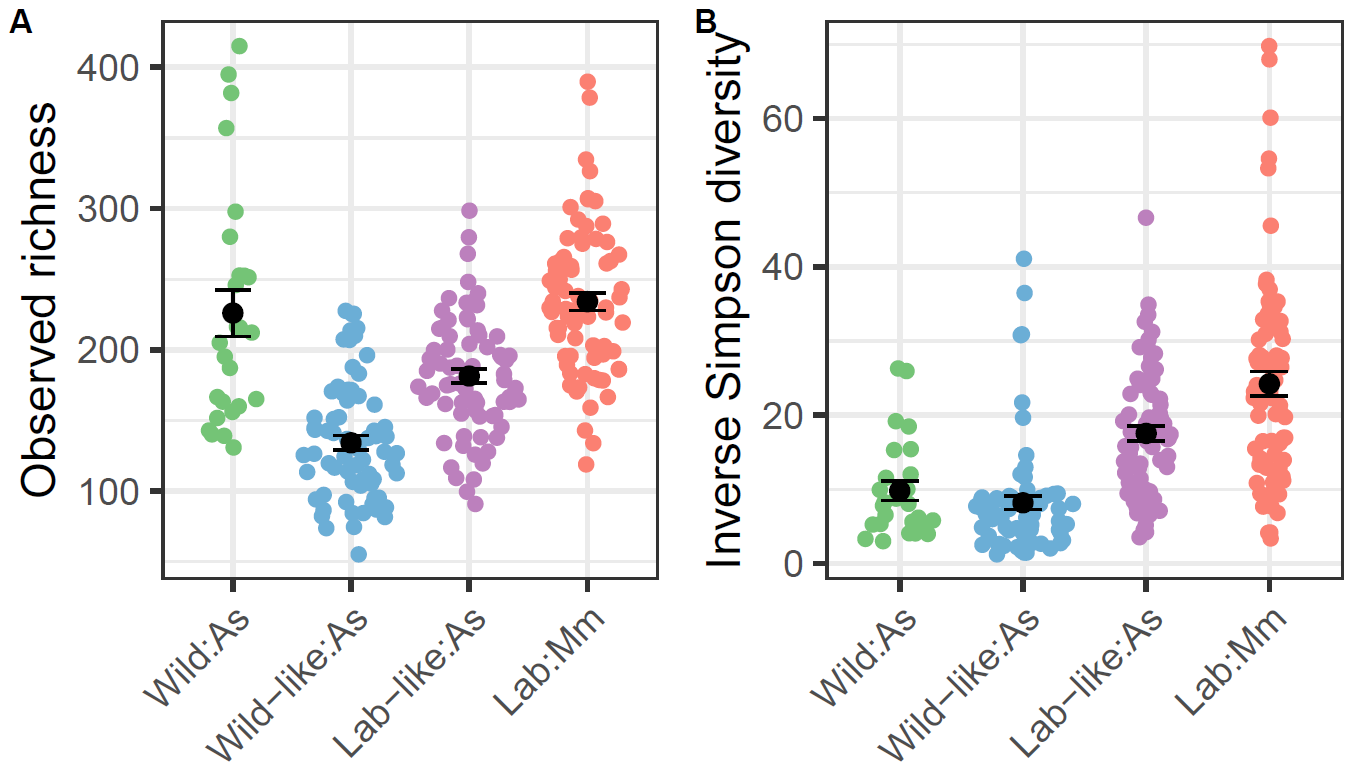

**Figure S2.** **Other gut microbial alpha diversity metrics among cohorts**: observed ASV richness **(A)** and inverse Simpson diversity **(B)** were calculated as the mean across 100 rarefactions to 15,857 reads per sample. Points represent individual samples; black points and error bars indicate group means ± SE.

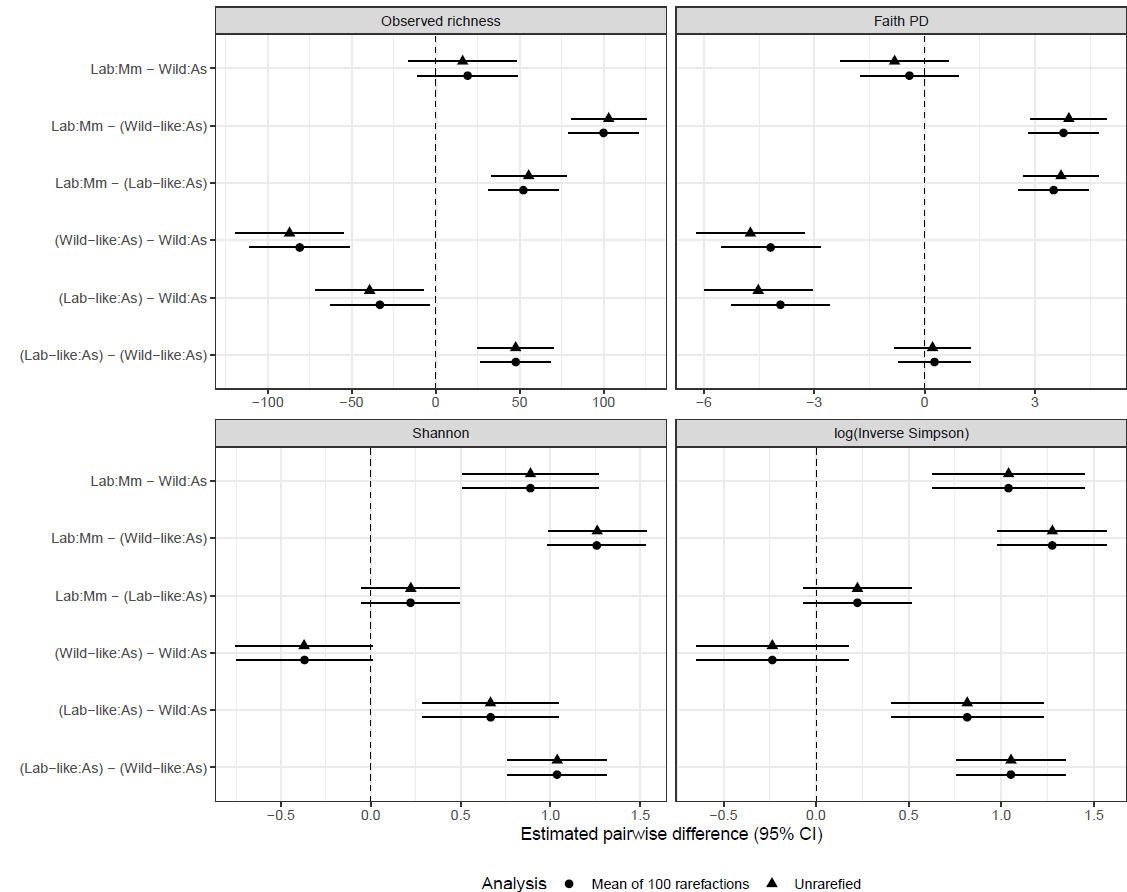

**Figure S3. Sensitivity of alpha-diversity results to rarefaction.** Pairwise differences among cohorts estimated using alpha-diversity values averaged across 100 rarefactions to 15,857 reads per sample (circles) and from unrarefied counts (triangles). Points show estimated marginal mean differences with 95% confidence intervals.

| **Table S2. Model results for four alpha diversity metrics.** | | | | |
| --- | --- | --- | --- | --- |
| **Alpha Diversity Metric** | **Predictor** | **Χ^2^** | **df** | **p value** |
| Observed richness | Cohort | 161.410 | 3 | <0.0001 |
| Observed richness | Sex | 1.093 | 1 | 0.296 |
| Faith PD | Cohort | 161.917 | 3 | <0.0001 |
| Faith PD | Sex | 0.280 | 1 | 0.597 |
| Shannon | Cohort | 167.237 | 3 | <0.0001 |
| Shannon | Sex | 0.122 | 1 | 0.727 |
| log(Inverse Simpson) | Cohort | 154.032 | 3 | <0.0001 |
| log(Inverse Simpson) | Sex | 0.169 | 1 | 0.681 |

| **Table S3. Pairwise comparisons of alpha diversity among cohorts.** Tukey-adjusted pairwise comparisons of estimated marginal means from models including cohort and sex as fixed effects and mouse identity as a random effect. Alpha-diversity estimates were averaged across 100 rarefactions to 15,857 reads per sample. | | | | | | | |
| --- | --- | --- | --- | --- | --- | --- | --- |
| **Metric** | **Cohort Contrast** | **Estimate** | **SE** | **95% LCI** | **95% UCI** | **Z ratio** | **P value** |
| Observed Richness | Lab:*Mm* - (Lab-like:*As*) | 52.265 | 8.061 | 31.555 | 72.974 | 6.484 | <0.0001 |
|  | Lab:*Mm* - (Wild-like:*As*) | 100.005 | 8.098 | 79.201 | 120.808 | 12.349 | <0.0001 |
|  | Lab:*Mm* - Wild:*As* | 19.089 | 11.545 | -10.569 | 48.748 | 1.654 | 0.349 |
|  | (Lab-like:*As*) - (Wild-like:*As*) | 47.740 | 8.128 | 26.859 | 68.621 | 5.874 | <0.0001 |
|  | (Lab-like:*As*) - Wild:*As* | -33.175 | 11.565 | -62.886 | -3.465 | -2.869 | 0.021 |
|  | (Wild-like:*As*) - Wild:*As* | -80.915 | 11.586 | -110.679 | -51.151 | -6.984 | <0.0001 |
| Faith PD | Lab:*Mm* - (Lab-like:*As*) | 3.503 | 0.373 | 2.544 | 4.463 | 9.381 | <0.0001 |
|  | Lab:*Mm* - (Wild-like:*As*) | 3.770 | 0.375 | 2.807 | 4.734 | 10.054 | <0.0001 |
|  | Lab:*Mm* - Wild:*As* | -0.414 | 0.522 | -1.755 | 0.926 | -0.794 | 0.857 |
|  | (Lab-like:*As*) - (Wild-like:*As*) | 0.267 | 0.376 | -0.700 | 1.234 | 0.710 | 0.893 |
|  | (Lab-like:*As*) - Wild:*As* | -3.918 | 0.523 | -5.260 | -2.575 | -7.497 | <0.0001 |
|  | (Wild-like:*As*) - Wild:*As* | -4.185 | 0.524 | -5.531 | -2.839 | -7.988 | <0.0001 |
| Shannon | Lab:*Mm* - (Lab-like:*As*) | 0.221 | 0.106 | -0.051 | 0.493 | 2.087 | 0.157 |
|  | Lab:*Mm* - (Wild-like:*As*) | 1.257 | 0.106 | 0.984 | 1.531 | 11.817 | <0.0001 |
|  | Lab:*Mm* - Wild:*As* | 0.888 | 0.148 | 0.509 | 1.267 | 6.019 | <0.0001 |
|  | (Lab-like:*As*) - (Wild-like:*As*) | 1.036 | 0.107 | 0.762 | 1.311 | 9.707 | <0.0001 |
|  | (Lab-like:*As*) - Wild:*As* | 0.667 | 0.148 | 0.287 | 1.047 | 4.513 | <0.0001 |
|  | (Wild-like:*As*) - Wild:*As* | -0.369 | 0.148 | -0.750 | 0.011 | -2.493 | 0.061 |
| log(Inverse Simpson) | Lab:*Mm* - (Lab-like:*As*) | 0.223 | 0.114 | -0.071 | 0.517 | 1.952 | 0.206 |
|  | Lab:*Mm* - (Wild-like:*As*) | 1.275 | 0.115 | 0.980 | 1.570 | 11.110 | <0.0001 |
|  | Lab:*Mm* - Wild:*As* | 1.039 | 0.159 | 0.629 | 1.449 | 6.517 | <0.0001 |
|  | (Lab-like:*As*) - (Wild-like:*As*) | 1.052 | 0.115 | 0.756 | 1.348 | 9.134 | <0.0001 |
|  | (Lab-like:*As*) - Wild:*As* | 0.816 | 0.160 | 0.405 | 1.226 | 5.107 | <0.0001 |
|  | (Wild-like:*As*) - Wild:*As* | -0.236 | 0.160 | -0.647 | 0.175 | -1.477 | 0.452 |

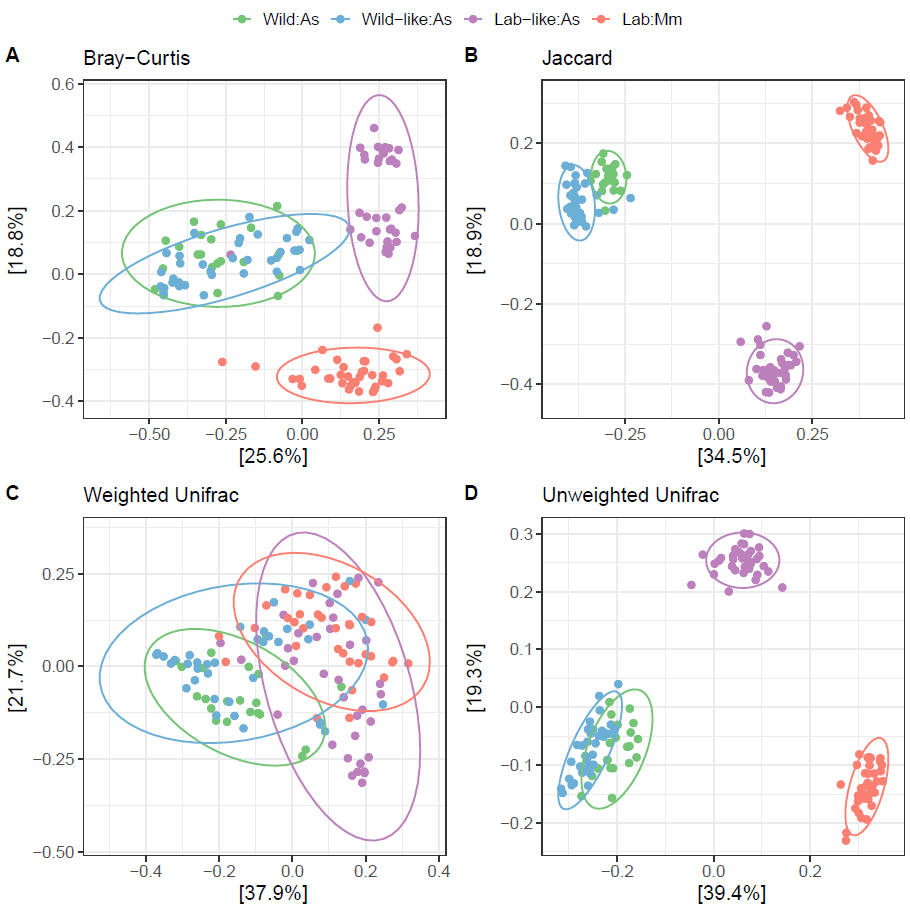

**Figure S4. Beta-diversity patterns among cohorts across four distance metrics.** PCoA of gut microbial community composition based on **(A)** Bray–Curtis dissimilarity, **(B)** Jaccard distance, **(C)** weighted UniFrac, and **(D)** unweighted UniFrac. Points represent individual mice, colours indicate cohort, and ellipses show the distribution of samples within each cohort. Axis labels indicate the percentage of variation explained by each principal coordinate. Bray-Curtis reflects abundance-weighted compositional differences, Jaccard reflects presence–absence differences, and weighted and unweighted UniFrac additionally incorporate phylogenetic relationships among ASVs.

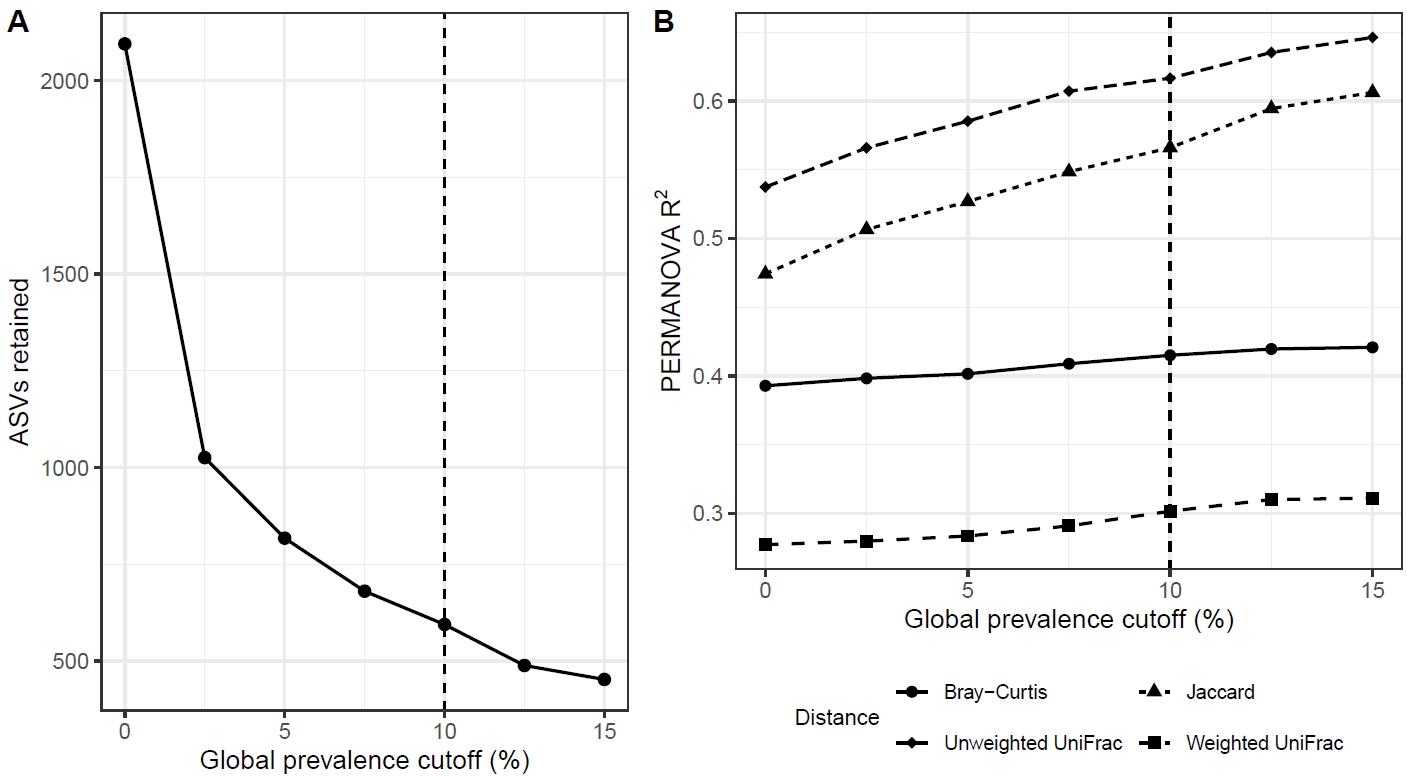

**Figure S5. Sensitivity of beta-diversity results to ASV prevalence filtering. (A)** Number of ASVs retained across global prevalence thresholds from 0% to 15%, with the dashed line indicating the 10% cutoff used in the primary analysis. (B) Overall PERMANOVA effect size (R^2^) for the effect of cohort on beta diversity across the same prevalence thresholds. Bray–Curtis and Weighted Unifrac effect sizes remained comparatively stable across filtering stringency, whereas Jaccard and Unweighted Unifrac effect sizes increased slightly as low-prevalence ASVs were removed.

| **Table S4. Overall PERMANOVA results for beta diversity.** | | | | | |
| --- | --- | --- | --- | --- | --- |
| **Distance metric** | **Fixed effect** | **df** | **pseudo-*F*** | ***R*²** | ***P*** |
| Bray–Curtis | Mouse cohort | 3 | 29.610 | 0.415 | <0.001 |
| Bray–Curtis | Sex | 1 | 1.018 | 0.005 | 0.390 |
| Jaccard | Mouse cohort | 3 | 56.897 | 0.576 | <0.001 |
| Jaccard | Sex | 1 | 1.368 | 0.005 | 0.180 |
| Weighted UniFrac | Mouse cohort | 3 | 18.707 | 0.310 | <0.001 |
| Weighted UniFrac | Sex | 1 | 0.810 | 0.004 | 0.518 |
| Unweighted UniFrac | Mouse cohort | 3 | 71.468 | 0.630 | <0.001 |
| Unweighted UniFrac | Sex | 1 | 1.591 | 0.005 | 0.130 |

| **Table S5. Pairwise PERMANOVA comparisons of gut microbial beta diversity among cohorts.** *P*-values were adjusted using the Benjamini–Hochberg procedure. | | | | |
| --- | --- | --- | --- | --- |
| **Distance metric** | **Comparison** | **pseudo-*F*** | ***R*²** | **BH-adjusted *P*** |
| **Bray–Curtis** | Wild:*As* vs Wild-like:*As* | 9.101 | 0.141 | <0.001 |
| **Bray–Curtis** | Wild:*As* vs Lab-like:*As* | 27.616 | 0.337 | <0.001 |
| **Bray–Curtis** | Wild:*As* vs Lab:*Mm* | 32.499 | 0.366 | <0.001 |
| **Bray–Curtis** | Wild-like:*As* vs Lab-like:*As* | 32.980 | 0.315 | <0.001 |
| **Bray–Curtis** | Wild-like:*As* vs Lab:*Mm* | 36.670 | 0.344 | <0.001 |
| **Bray–Curtis** | Lab-like:*As* vs Lab:*Mm* | 34.758 | 0.332 | <0.001 |
| **Jaccard** | Wild:*As* vs Wild-like:*As* | 13.653 | 0.198 | <0.001 |
| **Jaccard** | Wild:*As* vs Lab-like:*As* | 41.855 | 0.431 | <0.001 |
| **Jaccard** | Wild:*As* vs Lab:*Mm* | 55.809 | 0.502 | <0.001 |
| **Jaccard** | Wild-like:*As* vs Lab-like:*As* | 60.158 | 0.460 | <0.001 |
| **Jaccard** | Wild-like:*As* vs Lab:*Mm* | 91.102 | 0.564 | <0.001 |
| **Jaccard** | Lab-like:*As* vs Lab:*Mm* | 68.067 | 0.493 | <0.001 |
| **Weighted UniFrac** | Wild:*As* vs Wild-like:*As* | 7.311 | 0.116 | <0.001 |
| **Weighted UniFrac** | Wild:*As* vs Lab-like:*As* | 18.876 | 0.258 | <0.001 |
| **Weighted UniFrac** | Wild:*As* vs Lab:*Mm* | 24.410 | 0.305 | <0.001 |
| **Weighted UniFrac** | Wild-like:*As* vs Lab-like:*As* | 20.536 | 0.225 | <0.001 |
| **Weighted UniFrac** | Wild-like:*As* vs Lab:*Mm* | 23.606 | 0.253 | <0.001 |
| **Weighted UniFrac** | Lab-like:*As* vs Lab:*Mm* | 12.417 | 0.150 | <0.001 |
| **Unweighted UniFrac** | Wild:*As* vs Wild-like:*As* | 16.906 | 0.233 | <0.001 |
| **Unweighted UniFrac** | Wild:*As* vs Lab-like:*As* | 42.259 | 0.432 | <0.001 |
| **Unweighted UniFrac** | Wild:*As* vs Lab:*Mm* | 74.610 | 0.574 | <0.001 |
| **Unweighted UniFrac** | Wild-like:*As* vs Lab-like:*As* | 67.968 | 0.489 | <0.001 |
| **Unweighted UniFrac** | Wild-like:*As* vs Lab:*Mm* | 125.378 | 0.639 | <0.001 |
| **Unweighted UniFrac** | Lab-like:*As* vs Lab:*Mm* | 89.501 | 0.560 | <0.001 |

| **Table S6. Tests of homogeneity of multivariate dispersion among mouse cohorts.** PERMDISP assesses whether within-group dispersion differed among cohorts for each beta-diversity distance metric. | | |
| --- | --- | --- |
| **Distance metric** | ***F*** | ***P*** |
| **Bray–Curtis** | **1.367** | **0.261** |
| **Jaccard** | **18.289** | **<0.001** |
| **Weighted UniFrac** | **0.987** | **0.404** |
| **Unweighted UniFrac** | **16.119** | **<0.001** |

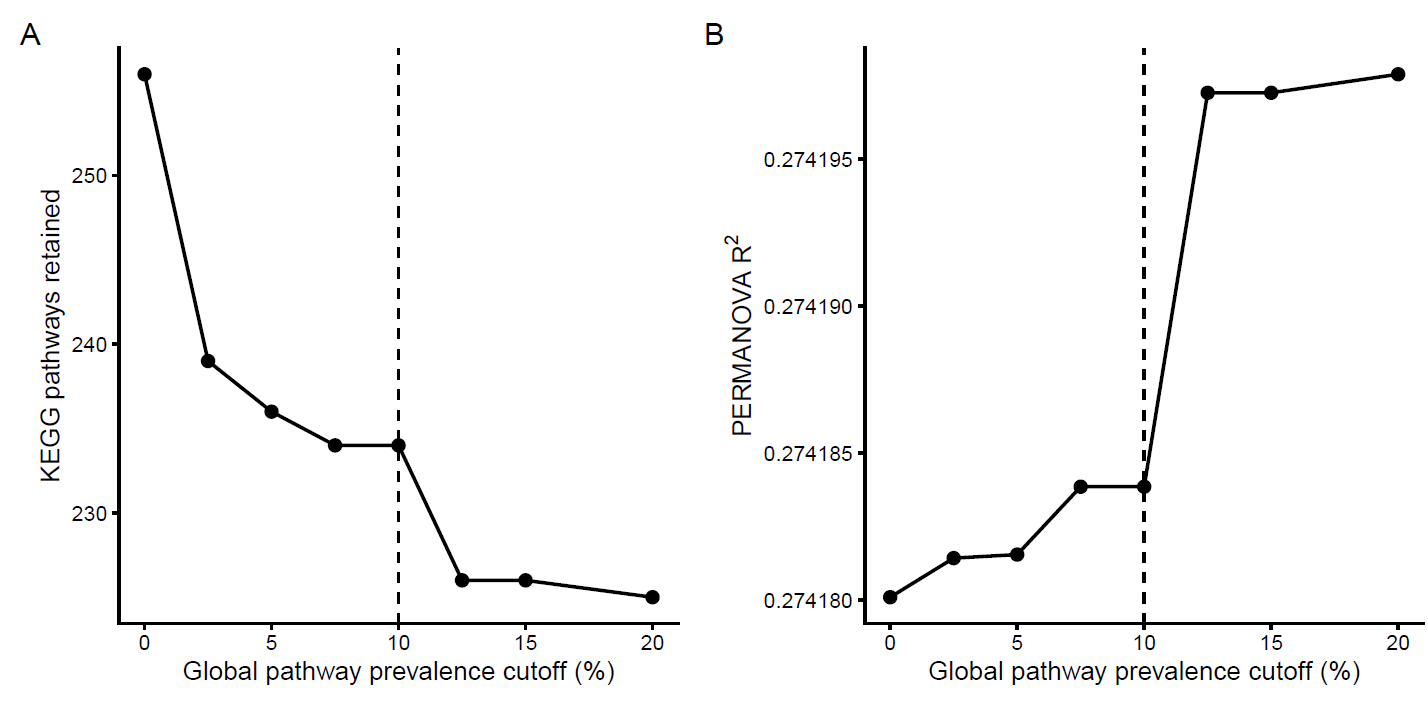

**Figure S6. Sensitivity of predicted KEGG functional composition to pathway-prevalence filtering.** Predicted KEGG pathways were filtered across global prevalence thresholds ranging from 0% to 20%. (A) Number of KEGG pathways retained at each prevalence threshold. (B) PERMANOVA (R^2^) for the effect of cohort on Bray–Curtis dissimilarity of predicted pathway composition at each threshold. The dashed vertical line indicates the 10% prevalence cutoff used in the primary analysis. Although increasingly stringent filtering reduced the number of retained pathways, the estimated cohort effect remained effectively unchanged across thresholds, indicating that the functional beta-diversity result was robust to the prevalence-filtering decision.

| **Table S6. Weighted nearest-sequenced-taxon index (NSTI) values for PICRUSt2 predictions across cohorts.** Differences were tested using a linear mixed-effects model with mouse identity as a random effect. The overall effect of cohort was not significant (*F*₃,₁₃₇.₃ = 1.84, *P* = 0.144). | | | | | | |
| --- | --- | --- | --- | --- | --- | --- |
| **Cohort** | **n** | **Mean NSTI** | **SD** | **Median** | **Minimum** | **Maximum** |
| Wild:*As* | 26 | 0.0462 | 0.0074 | 0.0458 | 0.0342 | 0.0599 |
| Wild-like:*As* | 70 | 0.0457 | 0.0068 | 0.0452 | 0.0335 | 0.0741 |
| Lab-like:*As* | 71 | 0.0448 | 0.0059 | 0.0449 | 0.0292 | 0.0591 |
| Lab:*Mm* | 72 | 0.0476 | 0.0076 | 0.0474 | 0.0331 | 0.0847 |
